# Unification of miRNA and isomiR research: the mirGFF3 format and the mirtop API

**DOI:** 10.1101/505222

**Authors:** Thomas Desvignes, Phillipe Loher, Karen Eilbeck, Jeffery Ma, Gianvito Urgese, Bastian Fromm, Jason Sydes, Ernesto Aparicio-Puerta, Victor Barrera, Roderic Espín, Eric Londin, Aristeidis G. Telonis, Elisa Ficarra, Marc R. Friedländer, John H. Postlethwait, Isidore Rigoutsos, Michael Hackenberg, Ioannis S. Vlachos, Marc K. Halushka, Lorena Pantano

**Author notes:** Corresponding author: Lorena Pantano.

## Abstract

**Background:** MicroRNAs (miRNAs) are small RNA molecules (∼22 nucleotide long) involved in post-transcriptional gene regulation. Advances in high-throughput sequencing technologies led to the discovery of isomiRs, which are miRNA sequence variants. While many miRNA-seq analysis tools exist, a lack of consensus on miRNA/isomiR analyses exists, and the resulting diversity of output formats hinders accurate comparisons between tools and precludes data sharing and the development of common downstream analysis methods.

**Findings:** To overcome this situation, we present here a community-based project, miRTOP (miRNA Transcriptomic Open Project) working towards the optimization of miRNA analyses. The aim of miRTOP is to promote the development of downstream analysis tools that are compatible with any existing detection and quantification tool. Based on the existing GFF3 format, we first created a new standard format, mirGFF3, for the output of miRNA/isomiR detection and quantification results from small RNA-seq data. Additionally, we developed a command line Python tool, ‘mirtop’, to manage the mirGFF3 format. Currently, mirtop can convert into mirGFF3 the outputs of commonly used pipelines, such as seqbuster, miRge2.0, isomiR-SEA, sRNAbench, and *Prost!*, as well as BAM files. Its open architecture enables any tool or pipeline to output results in mirGFF3.

**Conclusions:** Collectively a comprehensive isomiR categorization system, along with the accompanying mirGFF3 and mirtop API provide a complete solution for the standardization of miRNA and isomiR analysis, enabling data sharing, reporting, comparative analyses, and benchmarking, while promoting the development of common miRNA methods focusing on downstream steps to miRNA detection, annotation, and quantification.

## Background

microRNAs (miRNAs) are the best known class of small RNAs and were discovered in the nematode worm *C. elegans* [1,2]. It was first reported that the gene *lin-4* generated a 22 nucleotides (nt) long RNA molecule that bound to the 3’-UTR of the *lin-14* gene transcript, regulating its expression during larval development [3]. miRNA genes are transcribed into a primary RNA (*pri-miRNA*) that is processed into a hairpin-like miRNA precursor (*pre-miRNA*) after cutting off the 5’ and 3’-tails by Drosha and DGCR8 proteins [4]. The pre-miRNA hairpin is then exported to the cytoplasm and processed by Dicer, which cleaves off the hairpin loop and releases a miRNA duplex about 22 nt long [5]. Originally, it was believed that only one strand of the duplex is retained and incorporated into the RNA-induced silencing complex (RISC) thereby mediating gene silencing by imperfect base pairing between the miRNA and the 3’-UTR of target messenger RNAs (mRNA) [6]. It was later shown that miRNAs have a more expansive spectrum of activities. In fact, it was shown experimentally that both amino-coding sequences [7] and 5′-UTRs [8] can also be targeted by miRNAs. In addition, both arms of a miRNA can produce mature miRNAs, either simultaneously [9], or in a tissue specific manner [10,11]. miRNAs are essential to virtually all biological processes including, but not limited to, cell differentiation, cell proliferation, cell death, fat metabolism, and neuronal cell fate [1,2]. Moreover, the deficit or excess of miRNAs have been associated with several human diseases, such as, myocardial infarction and different types of cancer [12]. miRNAs reside not only inside cells, but also in a variety of biofluids [13–15], which has suggested they could be used as non-invasive disease biomarkers or therapies [16,17,18, 10].

IsomiRs are sequence variants from annotated miRNAs [19–21]. IsomiRs were first described in detail by Morin et al [22] in human stem cell lines using next generation sequencing technologies. Sequence variations can affect different parts of the mature miRNA sequence as consequences of different biochemical processes [23]. Variations at 5’ and 3’-ends could be due to imprecision of the Drosha/Dicer cutting machinery. However, it has already been shown that, in humans, these endpoint variations are constitutive in both healthy individuals and patients [10,24,25]. In fact, isomiRs have been shown to depend on a person’s sex, population origin, and ethnicity [10,24], as well as on tissue, tissue state, and disease subtype [26,27]. Non-templated nucleotide additions at the 3’-end can be due to nucleotide transferases (TUTase) that generally add adenine or uridine nucleotides [28]. 3′-end variations and non-templated additions take a new meaning considering that the tail end of a miRNA can contain a cell-compartment localization signal [29]. Finally, post-transcriptional processing of miRNAs can generate nucleotide changes at any position of the sequence by RNA processing enzymes, such as A-to-I editing by ADAR enzymes [30]. The specific function of isomiRs is still not well understood, but multiple studies have suggested a context-specific effect of isomiRs on gene regulation [26,28,31,32]. This is further supported by the fact that different isomiRs from the same mature miRNA can target virtually non-overlapping sets of transcripts [10,33–35].

Several tools have been developed to analyze miRNAs and their respective isomiRs [36] (Table 1). These tools differ in their alignment strategies, ways to handle cross-mapping events, abundance cutoffs, or isomiR annotation methods. All of them report isomiRs in a different format file and with different levels of complexity (Supp. File 1).

**Table 1.**
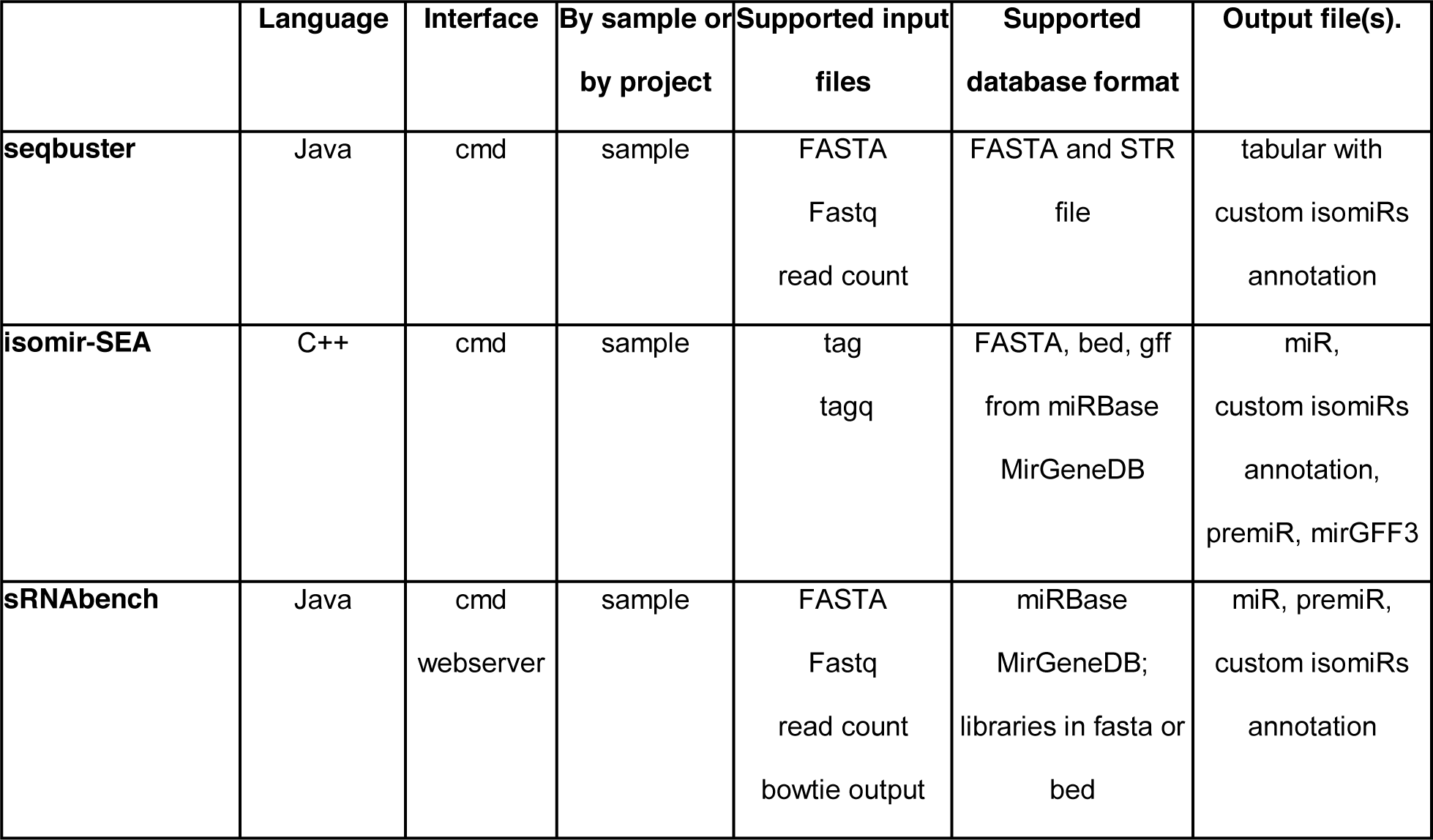
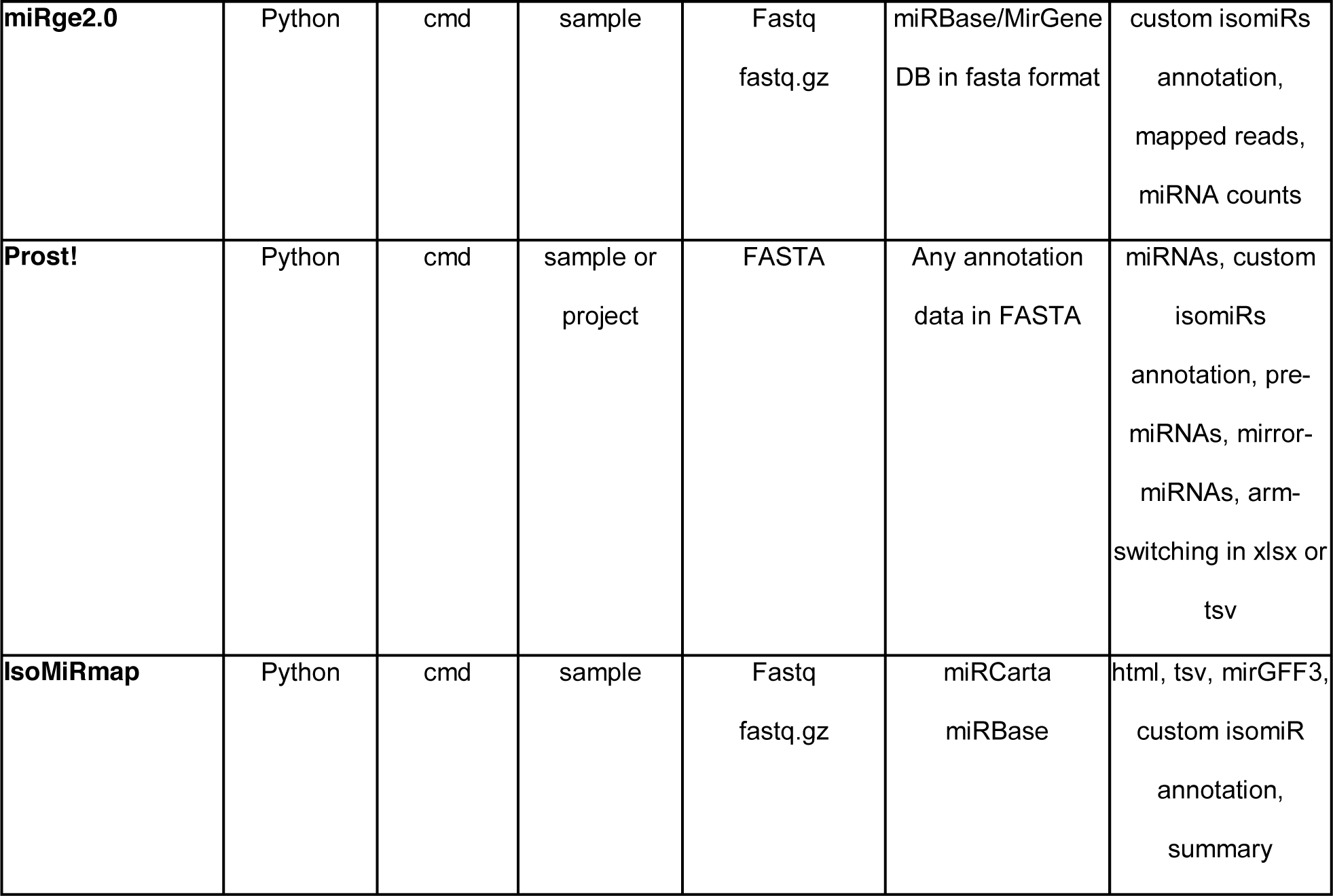
Overview of several tools reporting isomiR information from small RNA-seq data.

|  | Language | Interface | By sample or<br>by project | Supported input<br>files | Supported<br>database format | Output file(s). |
| --- | --- | --- | --- | --- | --- | --- |
| **seqbuster** | Java | cmd | sample | FASTA<br>Fastq<br>read count | FASTA and STR<br>file | tabular with<br>custom isomiRs<br>annotation |
| **isomir-SEA** | C++ | cmd | sample | tag<br>tagq | FASTA, bed, gff<br>from miRBase<br>MirGeneDB | miR,<br>custom isomiRs<br>annotation,<br>premiR, mirGFF3 |
| **sRNAbench** | Java | cmd<br>webserver | sample | FASTA<br>Fastq<br>read count<br>bowtie output | miRBase<br>MirGeneDB;<br>libraries in fasta or<br>bed | miR, premiR,<br>custom isomiRs<br>annotation |
| **miRge2.0** | Python | cmd | sample | Fastq<br>fastq.gz | miRBase/MirGene<br>DB in fasta format | custom isomiRs<br>annotation,<br>mapped reads,<br>miRNA counts |
| **Prost!** | Python | cmd | sample or<br>project | FASTA | Any annotation<br>data in FASTA | miRNAs, custom<br>isomiRs<br>annotation, pre-<br>miRNAs, mirror-<br>miRNAs, arm-<br>switching in xlsx or<br>tsv |
| **IsoMiRmap** | Python | cmd | sample | Fastq<br>fastq.gz | miRCarta<br>miRBase | html, tsv, mirGFF3,<br>custom isomiR<br>annotation,<br>summary |

Seqbuster integrates its own aligner to maximize the number of isomiRs analyzed but only retains isomiRs with a maximum of one nucleotide change within the miRNA and three changes at each end [23]. Seqbuster outputs a tabular delimited file with a column for each isomiR type. It works with the isomiR Bioconductor package to detect expression data and isomiR differences (https://doi.org/doi:10.18129/B9.bioc.isomiRs).

isomiR-SEA [37] implements a miRNA-specific alignment procedure for comparing each read of the sample to all the miRNA sequences from miRBase and MirGeneDB, collecting uniquely and multi-mapped sequences. The tool annotates the positions of the variations (mismatches and indels) enabling fine categorization of each aligned read that can be classified as canonical miRNA or one of the isomiRs described in Table 2. isomiR-SEA then outputs a detailed isomiR expression quantification table, with a focus on the conserved miRNA-mRNA interaction sites. isomiR-SEA is implemented in C++ using functions collected in the SeqAn bioinformatics library [38].

**Table 2.**
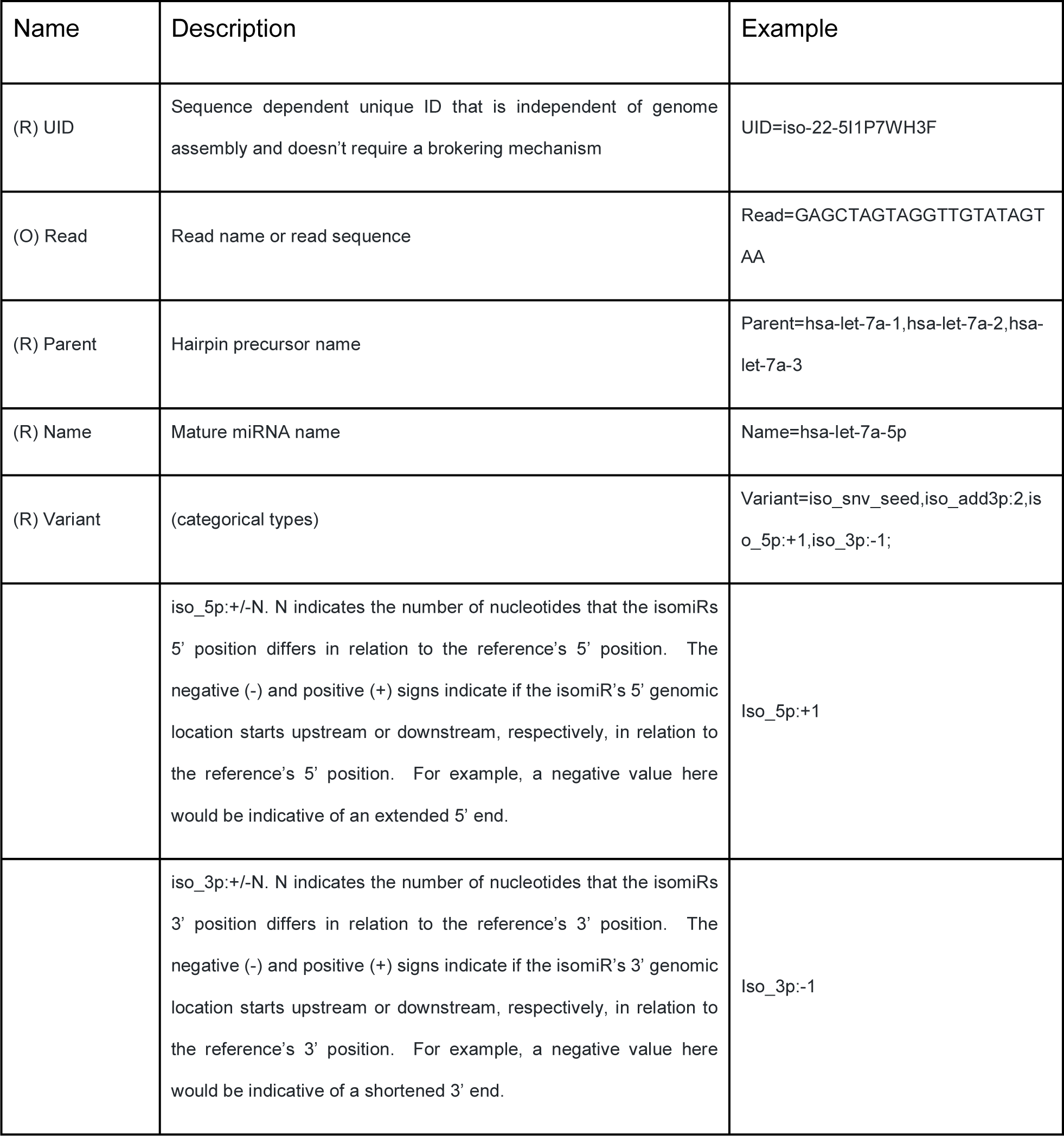
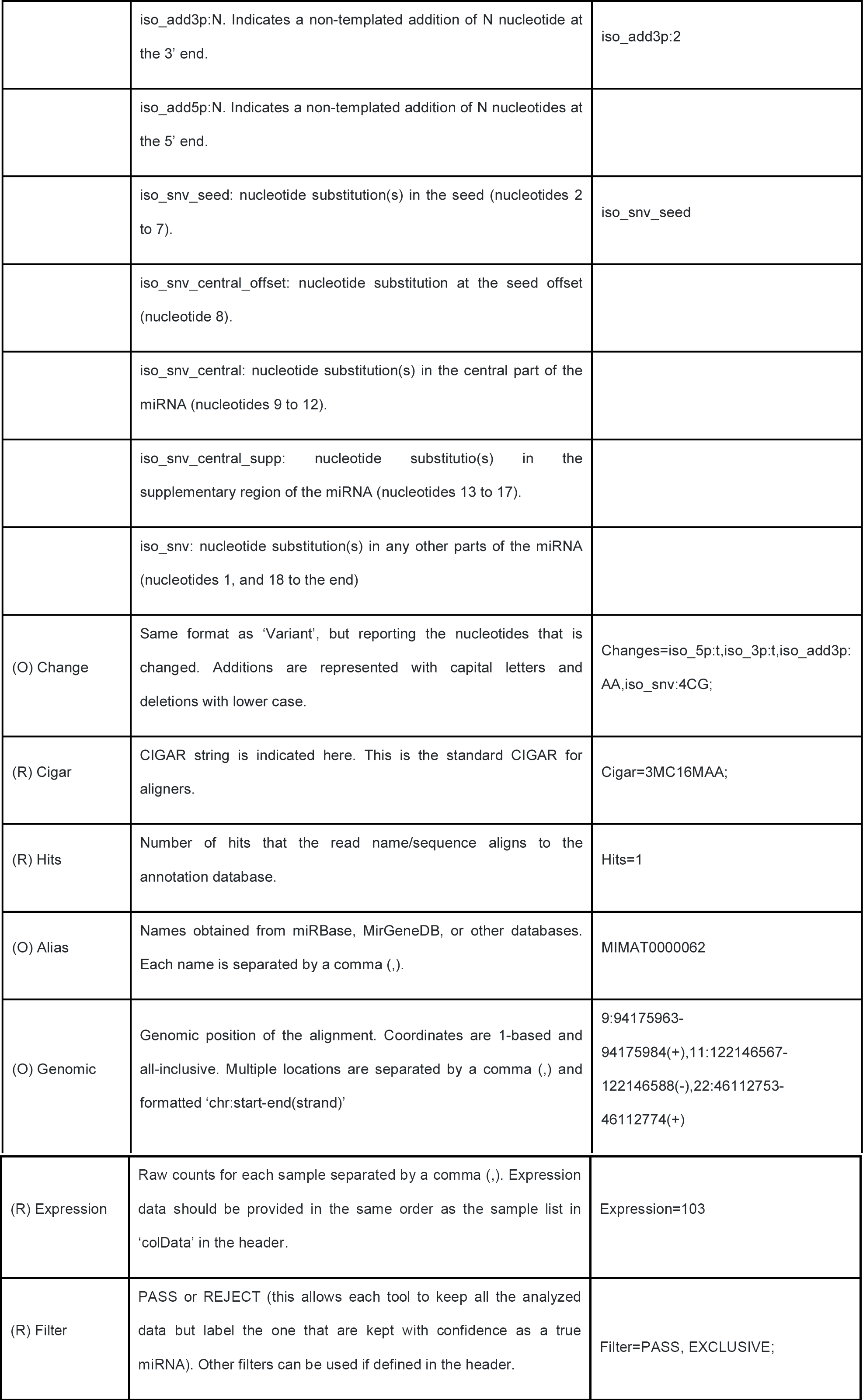
Description of the ‘Attribute’ column. Description of the attributes of the mirGFF3 format that can be stored in the ‘Attribute’ column (column 9). The first column shows the name of the attribute, and indicates whether it is a required (R) or an optional (O) value. The second column explains the nature of the attribute, and the third column provides an example based on an isomiR of the human miRNA hsa-let-7a-5p (also represented in Fig. 1C).

Another tool, sRNAbench [39,40], applies a bowtie seed alignment option, either to the genome (genome mode) or to miRNA reference sequences (library mode), to score only the first L nucleotides (by default L=19 allowing one mismatch) and therefore does not take into account mismatches at the 3’-end of the read caused by any post-transcriptionally added nucleotides. sRNAbench clusters all reads that map within a window of the canonical mature miRNA sequence (3 nt upstream of the start coordinate and 5 nt downstream of the end coordinate) and applies a hierarchical isomiR classification scheme. The sRNAbench tool has several tab-separated output files for isomiR analysis.

miRge2.0 maximizes isomiR discovery by iteratively mapping reads to user-defined miRNA and non-miRNA libraries using bowtie with a final step of loose alignment to the miRNA reads of any unaligned sequences [41]. miRge2.0, has a threshold option to remove called miRNAs whose reads are predominately isomiR, rather than canonical, based on a user-specified threshold to correct for false positive miRNAs.

*Prost!* (PRocessing Of Small Transcripts) quantifies and annotates miRNA expression [42]. *Prost!* uses the global aligner BBMap (https://sourceforge.net/projects/bbmap/) to align transcripts to a user-specifiable genome allowing for the identification of post-transcriptional modifications (e.g non-templated additions, editing, alternative cutting). *Prost!* then groups transcripts based on genomic location(s) and each group of sequences is annotated with user-defined databases of mature miRNAs, miRNA precursors, and other types of RNAs. Genomic location groups with identical annotations are further combined and can be used for downstream differential expression analyses.

IsoMiRmap maps and quantifies isomiRs by considering both miR-space and the rest of the genome. Many tools operate by mapping sequenced reads on a curated database such as miRBase or miRCarta. Some allow the user to provide her or his database of interest: e.g., all members of the let-7 family of miRNA precursors. The benefit of such an approach is its speedy execution due to the small size of the search space. The IsoMiRmap tool (Loher and Rigoutsos – Personal Communication), currently in development, makes it possible to consider the entire genome when mapping while having modest computational requirements. Considering the entire genome has the advantage of being able to flag whether or not an isomiR is exclusive to miR-space or if it could have been transcribed elsewhere. The IsoMiRmap tool ouputs in various formats, including HTML, tab separated files, and miRGFF3.

Although the analysis of miRNAs and their isomiRs has dramatically changed over the past several years, there is a lack of consensus among bioinformatic tools to annotate and study the isomiR landscape. Tools generate different output file types with different structures and isomiR notations. This lack of homogeneity prevents accurate comparisons between tools and precludes data sharing and the development of common downstream analyses which would be independent of the tool used for detection and quantification.

To overcome this situation, we present here mirGFF3, a standardized output format for the analysis of miRNAs and their isomiRs based on sequencing data. mirGFF3 was created to fit all research fields and as many tools as possible with the idea of democratization and standardization of miRNA analysis. This new file format allows the storage of relevant miRNA/isomiR information and was developed based on the existing GFF3 format [43], commonly used in genome annotation and messenger RNA analyses. Importantly, mirGFF3 uses an ontological strategy to relate the sequence features identified. Moreover, we developed a Python API - mirtop - that supports general file operations as well as importing miRNA tool output files and converting and exporting them into the new mirGFF3 format to promote the development of downstream tools usable by all in a collaborative environment.

## Findings

### The miRTop project

To communicate ideas, define standards, and to develop successful formats or tools that would be useful to the majority of researchers in the miRNA community, we created the miRTop community project (miRNA Transcriptomic Open Project), an entirely open source project. The project is open for participation to any member of the miRNA community, regardless of the level of seniority and status. miRTop serves as an incubator of ideas that helps improve miRNA analysis standards and boost collaboration. All updates and progress reports are and will continue to be publicly available as they happen and discussion summaries have already been released through GitHub. The project owns its own organization in GitHub articulated around four different repositories: 1) the main web page, 2) the mirGFF3 format, 3) the mirtop API, and 4) the incubator [44], where new ideas take form. Additional repositories will be added as projects develop. The miRTop group uses the GitHub project web pages to organize the different analyses in a transparent, communicative, and inclusive manner to promote collaboration and equality among all members of the miRNA community. Crowd-supported projects have recently started to emerge in bioinformatics research [45]. miRTop is a true trailblazer, enabling communication, collaboration, and community-driver problem solving and decision making. The research problems are selected by the miRNA community and commonly addressed.

### The mirGFF3 file format: definition and explanation

The mirGFF3 format was developed based on the original GFF3 format, taking advantage of the coordinate system information it can handle and the possibility to store attributes in column 9 (Supp. File 2). GFF3 format is commonly used for the annotation of genomic coordinates and is a popular data exchange format, particularly within Generic Model Organism Database (GMOD) [46] and genome browsing applications such as Ensembl or IGV. mirGFF3 format definition and corresponding descriptions are maintained on the mirGFF3 specific GitHub page, and have been deposited in the FAIRsharing [47] and EDAM databases [48].

The columns ‘seqid’, ‘source’, ‘type’, ‘start’, ‘end’, and ‘strand’, are used as defined in the original GFF3 format. The column ‘score’ is available for each tool to use freely if additional information needs to be added or specified. The column ‘phase’ is ignored in the mirGFF3 format given that it refers specifically to coding sequences. Finally, the column 9, ‘attribute’, was adapted to contain all the relevant information concerning the metadata that characterize each specific isomiR (Table 2). In the mirGFF3 definition, attributes starting with a capitalized letter are reserved to the attributes listed in Table 2, but custom attributes can be added by adding their descriptor in lower case.

mirGFF3 format accepts headers that include sample origin, names, and other custom information used to parse the data by the API framework. All header lines should start with the string `##`. Only three lines are mandatory: the mirGFF3 format version, the database used for annotation, and the sample name (Figure 1A). The database line can point to any of the already published resources: miRBase [49], miRCarta [50], miRGeneDB [51], or a custom database. For the existing databases, the version should be provided. For the custom databases, an optional link to download the coordinates or precursor sequences would be the desired information to add. Sample names should be given after the character string `COLDATA` and should contain the sample names, each separated by a `,` (comma) character. If the attribute ‘Filter’ is used, a line starting with the character string `FILTER`, explaining the possible values this attribute refers to, should be added for the user to filter the file content based on these criteria (Figure 1A). In addition, we encourage users to add any header line that could provide additional useful information concerning the original small RNA-seq analysis. The most common information could be, for example, the annotation database used, the command line/parameters used to generate the mirGFF3 file, its date, or the description of any custom adjustments done during the previous annotation and quantification analysis.

**Figure 1.**
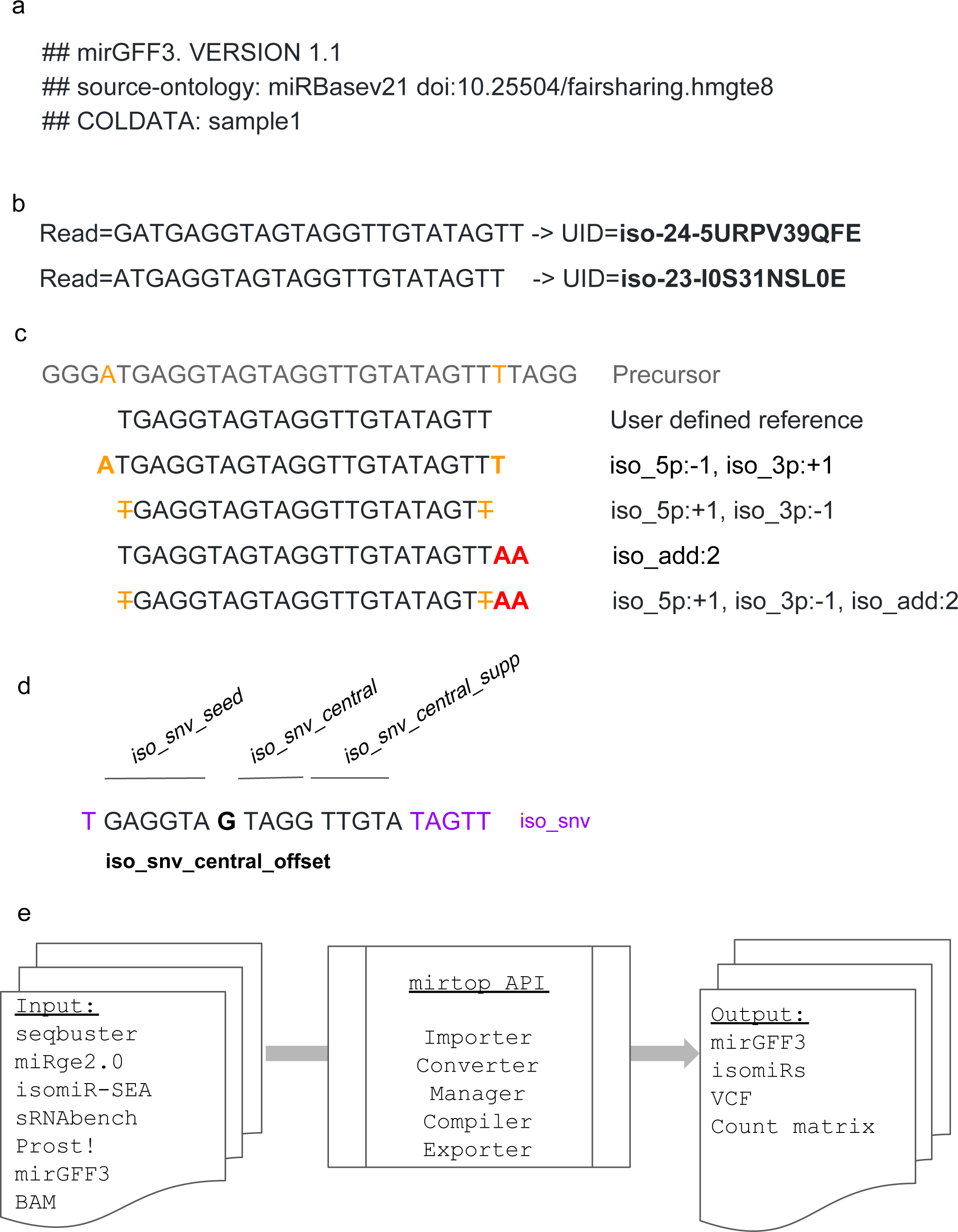
The mirGFF3 file format and the mirtop API. a) Example of input file header and the required lines: file format version, database used, and samples included. b) Examples of sequence compression to uniquely identify each sequence. c) Examples of isomiRs with changes at the 5’- and 3’-ends and their respective variant attributes. The first sequence represents a portion of the miRNA precursor (i.e. pre-miRNA) and the second sequence is the reference isomiR defined by the user or the database used. Bold orange letters indicate templated additions. Bold red letters indicate non-templated additions at the 3’-end of the isomiR. Orange strikethrough letters indicate non-templated nucleotide additions at the 3’-end of the isomiR. d) Example of nucleotide changes at different positions/regions of the isomiR and their respective naming. e) The mirtop API workflow shows the main formats accepted as input files, the functions the Python API have implemented, and the output file formats.

The column ‘attribute’ (column 9) of the original GFF3 format was adapted to contain all the metadata relevant to miRNA/isomiR analyses (e.g. the sequence variation(s) that an isomiR displays, its expression in different samples) with the possibility to filter and classify isomiRs by mature miRNA(s) and/or precursor name(s) (Table 2). The proper identification of isomiRs between studies is still an open problem, since different tools and research groups utilize different nomenclatures and representations. To this end, we adopted a unique identifier (UID), or ‘IsomiR license plate’, inspired by the MINTmap approach for tRNA fragments [52,53] and acting as a sequence-dependent unique ID that is independent of genome assembly and doesn’t require a brokering naming mechanism. Any isomiR sequence can be mapped to a UID and any UID can be converted back to the isomiR sequence it represents (Figure 1B).

**Table 2:** Description of the mirGFF3 ‘Attribute’ column (column 9).

Another characteristic of the mirGFF3 format which is specifically devised to maximize clarity, communication and standardization across the community is the *‘Variant’* attribute, which follows the isomiR description and miRNA-mRNA interaction sites adapted from isomiR-SEA format [37]. Briefly, modifications are based on comparing the sequence of a given isomiR to the reference miRNA in the chosen database. Changes on the 5’-end of the sequence, related to the start of the miRNA, are described as *‘iso_5p’* and changes on the 3’-end of the sequence, related to the tail of the miRNA, are described as *‘iso_3p’*. They are followed by the number of nucleotides that change in this isomiR compared to the defined reference. A ‘-’ (minus sign) is used if the isomiR startpoint is upstream compared to the reference startpoint. In contrast, a ‘+’ (plus sign) is used if the isomiR endpoint is downstream compared to the reference endpoint (Figure 1C) [10,24]. For example, an isomiR that has both ‘iso_5p:-1’ and ‘iso_3p:+1’, would be two nt longer than the reference: one nt longer on the 5’-end and one nt longer on the 3’-end (Figure 1C). In both *‘iso_5p’* and *‘iso_3p’* cases, the nucleotide additions have to be templated additions, meaning that these nucleotides are genomically encoded. In the case of 3’-non-templated additions (an addition that doesn’t match the genomic sequence), the isomiR is described as *‘iso_add3p’* (Figure 1C). Similarly, in the case of 5’-non-templated additions, the isomiR is described as ‘*iso_add5p*’. Finally, isomiRs that present nucleotide changes in their sequence that do not affect their ends are described as *‘iso_snv’* (single nucleotide variant). This type of isomiR is further divided into five subtypes (Figure 1D): 1) *‘iso_snv_seed’*, when the nucleotide variation is located in the seed of the detected isomiR between nucleotides 2 to 7; 2) *‘iso_snv_central_offset’*, when the nucleotide variation is located at the seed offset position, at nucleotide 8, a nucleotide that is relevant to the strength of the miRNA-mRNA interaction; 3) *‘iso_snv_central’*, when the nucleotide variation is located in the central part of the miRNA, between nucleotide 9 to 12, 4) *‘iso_snv_central_supp’*, when the nucleotide variation is located in the supplementary region of the miRNA, between nucleotides 13 to 17; and 5) *‘iso_snv’*, when the nucleotide variation is located in any other position in the miRNA, nucleotides 1, and 18 to the end of the miRNA.

The ‘*Filter’* attribute was adapted from the variant caller format file, where it is used to decide whether a variant passes or not the user defined filtering options. We followed the same idea enabling each tool to attribute to each isomiR a reliability score that can be any custom value defined in the additional header lines of the mirGFF3 file.

The ‘*Hits’* attribute is used to represent the number of times that the read name/sequence matches the database with different isomiR changes. For example, in the human genome assembly GRCh38 [54], iso-21-DV0Y6O6NB (with sequence AATGCACCTGGGCAAGGATTA) can be attributed to both MIMAT0002871&hsa-miR-500a-3p (ATGCACCTGGGCAAGGATTCTG) and MIMAT0004775&hsa-miR-502-3p (AATGCACCTGGGCAAGGATTCA) with different but presumably equally likely variations from the two references. In the first assignment case (hsa-miR-500a-3p), the 5’-end differs from the reference by one nucleotide, whereas in the second assignment case (hsa-miR-502-3p), the 3’-end differs from the reference by one nucleotide. By setting ‘Hits=2’ and representing the sequence in two lines (with ‘*Parent*’ attribute being one of reference in each line), both possible origins can be adequately captured. The ‘Expression’ value for the variant is set to the number of total reads for the sequence, and not a proportion of it, and the ‘*UID*’ attribute can be used to parse the file and avoid over-counting. A different example could be the isomiR iso-23-UPVMX5I80O (with sequence TACAGTAGTCTGCACATTGGTTA) that can be attributed to three different loci located on three different chromosomes: MIMAT0004563&hsa-miR-199b-3p on chromosome 9, and MIMAT0000232&hsa-miR-199a-3p on chromosomes 1 and 19 (all three loci having ACAGTAGTCTGCACATTGGTTA as reference sequence). In this situation, one can set “Hits=3” and take a similar approach as above. Alternatively, because this isomiR perfectly matches each genomic location, it could be listed in a single line with the ‘*Hits*’ attribute set to ‘1’ and the ‘*Parent*’ attribute would be used to reflect the multiple possible origin by having the three reference names separated by a comma character.

### The ‘mirtop’ API framework

The API framework ‘mirtop’ was developed in Python versions 2.7 and 3.6 and uses other common bioinformatics packages to support operations with BAM files (pysam) [55] and standard IO processes with sequences (Biopython) [56]. The mirtop package is based on a central class that converts each line of the mirGFF3 file into a Python class structure, containing all the information related to each isomiR. This method allows the centralization of every operation into a single file that can evolve over time without changing the functions that perform the operations. In addition, the validation of the file occurs simultaneously with the file being loaded, enabling the verification of the mirGFF3 rules and restrictions, avoiding errors that can be difficult to uncover later.

The mirtop API framework contains five different operations: importing, converting, managing, compiling, and exporting (Figure 1E). The importers in mirtop have been coded to import and convert the output files of seqbuster (bcbio-nextgen), miRge2.0, isomir-SEA, sRNAbench, and *Prost!* into the mirGFF3 format. Furthermore, IsoMiRmap, miRge2 and QuagmiR [57], have been extended by their groups to output results directly to mirGFF3 format. This is a clear indication of the shorter adaptation time required for new standards originating from community-driven projects. mirtop operator can manage and compile mirGFF3 files allowing joining, filtering on single or multiple files, and transformation of the mirGFF3 information into a count matrix. Finally, mirtop exporters create the final mirGFF3 file and can also convert it into other output formats commonly used for downstream analyses. Currently, mirtop, in addition to the mirGFF3 format, can export to isomiRs (Bioconductor package) and VCF formats, which are used in a diversity of visualization and analysis tools for isomiR characterization and variant calling (https://bioconductor.org/packages/release/bioc/html/isomiRs.html, http://www.internationalgenome.org/wiki/Analysis/vcf4.0/).

The conversion of several different tool outputs into a common file format will help researchers and developers focus on downstream analyses without being limited to only one quantifying tool and a specific output format. The mirtop API will therefore help spur the development of universal downstream analyses, enhancing the quality of miRNA and isomiR biology research.

## Conclusion

Here we present a community-backed effort to standardize, homogenize, and enhance the ways we report, share, and communicate miRNA results. We have organized as a community using miRTop as a common goal for miRNA/isomiR result standardization, and created mirGFF3, an adapted GFF3 file format. mirGFF3 was specifically designed to contain all relevant information concerning miRNAs and isomiRs identified in small RNA-seq data, regardless of the upstream methods or downstream use-cases. This new format represents the first consensus for isomiR variations and abundance report in one or more biological samples produced by high throughput sequencing technologies. The mirGFF3 format is complementary to existing bioinformatics tools that support GFF3 files and aligns to the transcriptomic communities that have based their messenger RNA annotations on GFF3 files. Similar to BAM or VCF file formats, mirGFF3 contains all the information necessary to re-analyze the data in the same way as when the raw output file from any analysis pipeline is available. The API framework, mirtop, enabling the conversion of miRNA quantification tool outputs and the processing of general statistics and count matrices, will serve as a catalyst for the use of the mirGFF3 format. The mirtop API supports any version of the mirGFF3 format and can convert older files to the latest version if needed.

The mirGFF3 file format and the mirtop API tool are the results of an open-membership international miRNA community created to promote open source code sharing in a collaborative and well-supported bioinformatic environment. The mirGFF3 format and associated mirtop API will encourage the miRNA community to develop common downstream analysis protocols, independent of the initial tool that was used for detection and quantification. The mirGFF3 format will provide a common entry point for a variety of applications ranging from the annotation of miRNAs/isomiRs or filtering for technical errors inherent to each library preparation protocol [58], to visualization, variant calling, differential expression, clustering, or any other sequence analyses.

The miRTOP group is and will remain open to any researcher interested in small RNA analysis at any level, from experimental scientists to computational biologists. miRTOP was created by members of the miRNA research community for the miRNA research community and offers networking and organization to improve and to promote collaborative research.

## Acknowledgments

Authors thank Peter Batzel for pointing us to the original GFF3 format, Rafael Alis for helping in the debugging the mirGFF3 conversion function, Yin Lu for integrating mirGFF3 into miRge2.0 and Shruthi Bandyadka for integrating the tabular exporter operation. T.D and J.H.P were supported by grant PLR-1543383 of the National Science Foundation, B.F. and M.R.F. acknowledge funding from the Strategic Research Area (SFO) program of the Swedish Research Council (VR) through Stockholm University, M.K.H. was supported by grant 1R01HL137811 of the National Institutes of Health, National Heart Lung Blood Institute, and was supported by the George and Marie Vergottis Fellowship of Harvard Medical School.

## Authors’ contributions

LP, TD, PL, KE, JS, BF, GU, EL, AGT, EF, MRF, JHP, IR, MH, MKH and ISV contributed to the conceptualization.

LP, TD, PL, BF, ISV and MKH contributed to the supervision of the project.

LP, SB, VB, RE, JM, PL, JS, GU and EAP contributed to the software development of mirtop or other tools to adapt to the mirGFF3 format.

LP, TD, GU, MH and MKH wrote the original draft.

All authors contributed to the review and editing of the final version.

## Availability and requirements

Use the *devel* branch in all repositories while reviewing this paper. We will update the *master* branch when the paper is published.

- Project name: miRTOP (miRNA Transcriptomic Open Project)
- Project home page: http://mirtop.github.io
- Source Code: https://github.com/miRTop/mirtop
- Operating system(s): Any
- Programming language: Python version 2.7 and 3.6
- Other requirements: pysam, pybedtools
- License: MIT

---

**Supplementary File 1:**
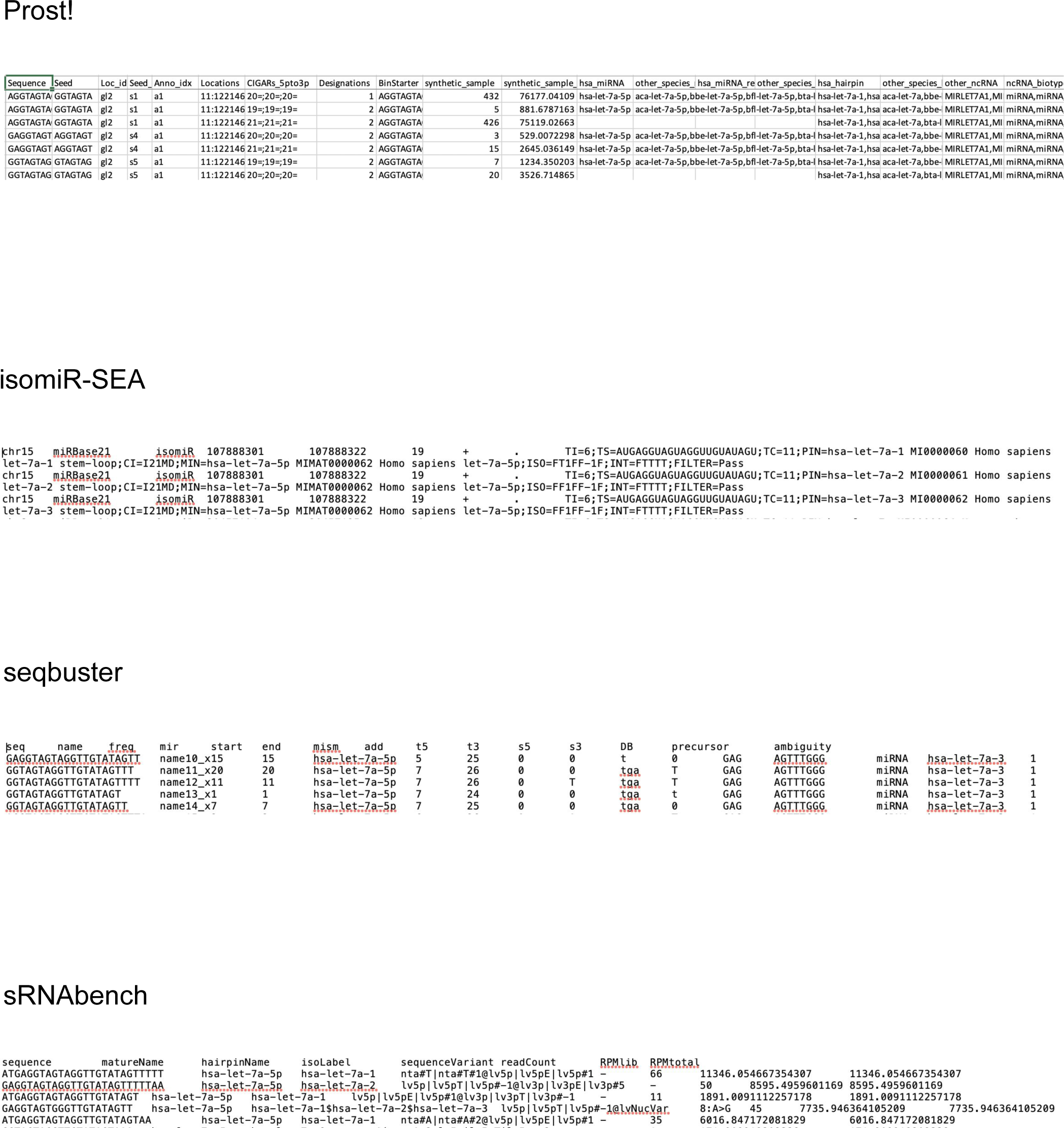
Example of output files from the tools described in Table 1. A few lines of the output of each tool is represented here to highlight the diversity of annotation and information given.

**Supplementary File 2:**
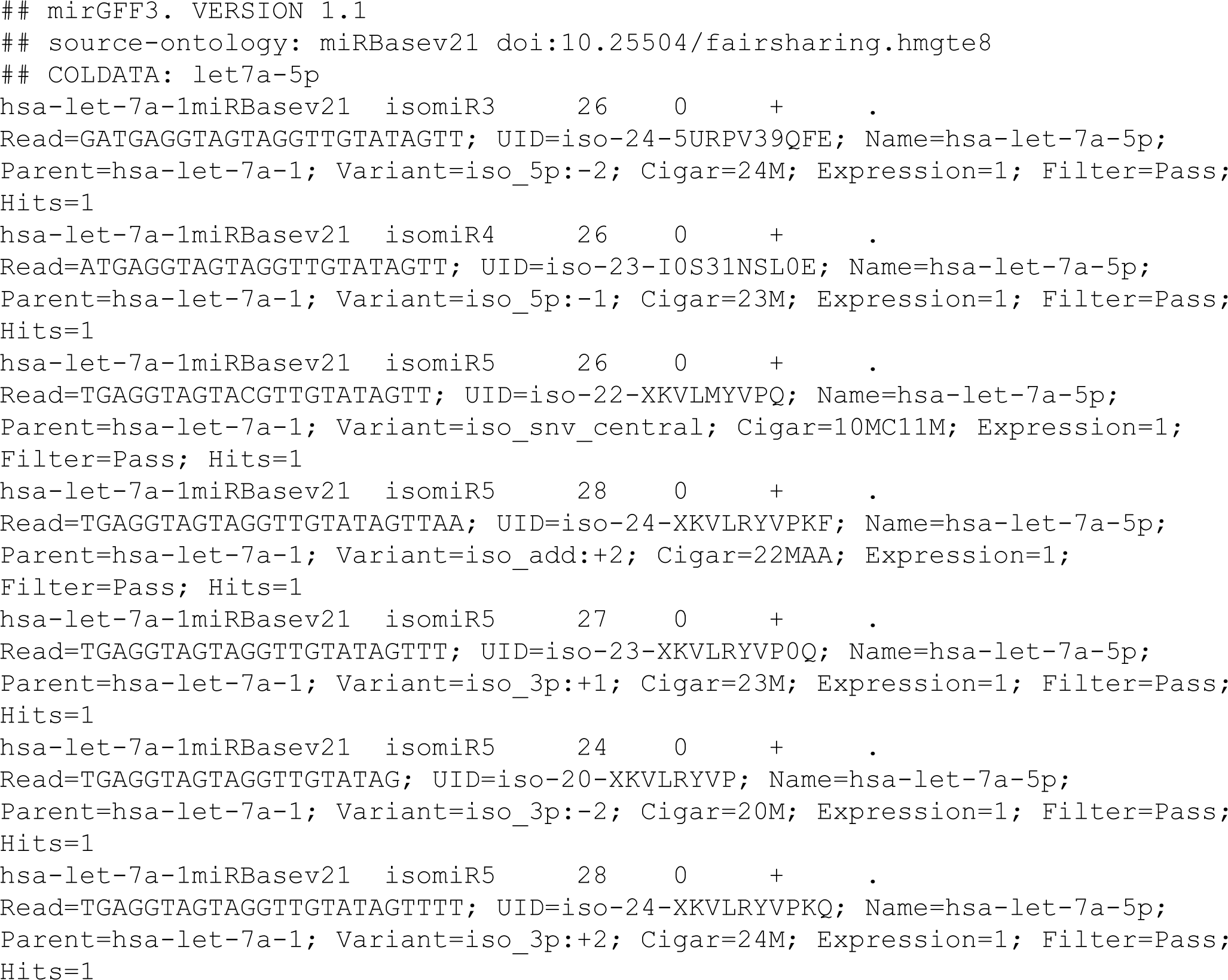
Example of a mirGFF3 file from simulated isomiRs of the hsa-let-7a miRNA.

## List of abbreviations

miRNA: microRNA
mRNA: messenger RNA
nt: nucleotide
RISC: RNA-induced silencing complex
UID: unique identifier
UTR: Untranslated region.

## Bibliography

1. Bartel DP. MicroRNAs: genomics, biogenesis, mechanism, and function. Cell. 2004;116: 281–297.

2. Bartel DP. Metazoan MicroRNAs. Cell. Elsevier; 2018;173: 20–51.

3. Lee RC, Feinbaum RL, Ambros V. The C. elegans heterochronic gene lin-4 encodes small RNAs with antisense complementarity to lin-14. Cell. 1993;75: 843–854.

4. Denli AM, Tops BBJ, Plasterk RHA, Ketting RF, Hannon GJ. Processing of primary microRNAs by the Microprocessor complex. Nature. 2004;432: 231–235.

5. Perron MP, Provost P. Protein interactions and complexes in human microRNA biogenesis and function. Front Biosci. 2008;13: 2537–2547.

6. Vella MC, Reinert K, Slack FJ. Architecture of a validated microRNA::target interaction. Chem Biol. 2004;11: 1619–1623.

7. Tay Y, Zhang J, Thomson AM, Lim B, Rigoutsos I. MicroRNAs to Nanog, Oct4 and Sox2 coding regions modulate embryonic stem cell differentiation. Nature. 2008;455: 1124–1128.

8. Zhou H, Rigoutsos I. MiR-103a-3p targets the 5′ UTR ofGPRC5Ain pancreatic cells. RNA. 2014;20: 1431–1439.

9. Yang J-S-S. Yang J, Phillips MD, Betel D, Mu P, Ventura A, et al. Widespread regulatory activity of vertebrate microRNA* species. RNA. 2010;17: 312–326.

10. Telonis AG, Loher P, Jing Y, Londin E, Rigoutsos I. Beyond the one-locus-one-miRNA paradigm: microRNA isoforms enable deeper insights into breast cancer heterogeneity. Nucleic Acids Res. 2015;43: 9158–9175.

11. Londin E, Loher P, Rigoutsos I. Reply to Backes and Keller: Identification of novel tissue-specific and primate-specific human microRNAs. Proc Natl Acad Sci U S A. 2015;112: E2851.

12. Ardekani AM, Naeini MM. The Role of MicroRNAs in Human Diseases. Avicenna J Med Biotechnol. 2010;2: 161–179.

13. Zhou X, Lu Z, Wang T, Huang Z, Zhu W, Miao Y. Plasma miRNAs in diagnosis and prognosis of pancreatic cancer: A miRNA expression analysis. Gene. 2018;673: 181–193.

14. Zhang Z, Xie Q, He D, Ling Y, Li Y, Li J, et al. Circular RNA: new star, new hope in cancer. BMC Cancer. 2018;18: 834.

15. Umu SU, Lyle R, Langseth H, Rounge TB. PO-096 Natural variation in serum small non-coding rnas – potential biomarkers of cancer. Poster Presentation: Prevention and Early Detection. 2018. doi:10.1136/esmoopen-2018-eacr25.624

16. Liu N, Cui R-X, Sun Y, Guo R, Mao Y-P, Tang L-L, et al. A four-miRNA signature identified from genome-wide serum miRNA profiling predicts survival in patients with nasopharyngeal carcinoma. Int J Cancer. 2014;134: 1359–1368.

17. Zhang Y, Roth JA, Yu H, Ye Y, Xie K, Zhao H, et al. A 5-MicroRNA Signature Identified from Serum MicroRNA Profiling Predicts Survival in Patients with Advanced Stage Non-Small Cell Lung Cancer. Carcinogenesis. 2018; doi:10.1093/carcin/bgy132

18. Pan J, Zhou C, Zhao X, He J, Tian H, Shen W, et al. A two-miRNA signature (miR-33a-5p and miR-128-3p) in whole blood as potential biomarker for early diagnosis of lung cancer. Sci Rep. 2018;8: 16699.

19. Kim H, Kim J, Kim K, Chang H, You K, Narry Kim V. AQ-seq: Accurate quantification of microRNAs and their variants [Internet]. 2018. doi:10.1101/339606

20. Fromm B, Billipp T, Peck LE, Johansen M, Tarver JE, King BL, et al. A Uniform System for the Annotation of Vertebrate microRNA Genes and the Evolution of the Human microRNAome. Annu Rev Genet. 2015;49: 213–242.

21. Desvignes T, Batzel P, Berezikov E, Eilbeck K, Eppig JT, McAndrews MS, et al. miRNA Nomenclature: A View Incorporating Genetic Origins, Biosynthetic Pathways, and Sequence Variants. Trends Genet. 2015;31: 613–626.

22. Morin RD, O’Connor MD, Griffith M, Kuchenbauer F, Delaney A, Prabhu A-L, et al. Application of massively parallel sequencing to microRNA profiling and discovery in human embryonic stem cells. Genome Res. 2008;18: 610–621.

23. Pantano L, Estivill X, Martí E. SeqBuster, a bioinformatic tool for the processing and analysis of small RNAs datasets, reveals ubiquitous miRNA modifications in human embryonic cells. Nucleic Acids Res. 2010;38: e34.

24. Loher P, Londin ER, Rigoutsos I. IsomiR expression profiles in human lymphoblastoid cell lines exhibit population and gender dependencies. Oncotarget. 2014;5: 8790–8802.

25. Telonis AG, Rigoutsos I. Race Disparities in the Contribution of miRNA Isoforms and tRNA-Derived Fragments to Triple-Negative Breast Cancer. Cancer Res. 2018;78: 1140–1154.

26. Telonis AG, Magee R, Loher P, Chervoneva I, Londin E, Rigoutsos I. Knowledge about the presence or absence of miRNA isoforms (isomiRs) can successfully discriminate amongst 32 TCGA cancer types. Nucleic Acids Res. 2017;45: 2973–2985.

27. Magee RG, Telonis AG, Loher P, Londin E, Rigoutsos I. Profiles of miRNA Isoforms and tRNA Fragments in Prostate Cancer. Sci Rep. 2018;8: 5314.

28. Menezes MR, Balzeau J, Hagan JP. 3’ RNA Uridylation in Epitranscriptomics, Gene Regulation, and Disease. Front Mol Biosci. 2018;5: 61.

29. Hwang H-W-W. Hwang H, Wentzel EA, Mendell JT. A Hexanucleotide Element Directs MicroRNA Nuclear Import. Science. 2007;315: 97–100.

30. Kawahara Y, Zinshteyn B, Sethupathy P, Iizasa H, Hatzigeorgiou AG, Nishikura K. Redirection of silencing targets by adenosine-to-inosine editing of miRNAs. Science. 2007;315: 1137–1140.

31. Garate X, La Greca A, Neiman G, Blüguermann C, Santín Velazque NL, Moro LN, et al. Identification of the miRNAome of early mesoderm progenitor cells and cardiomyocytes derived from human pluripotent stem cells. Sci Rep. 2018;8: 8072.

32. Trontti K, Väänänen J, Sipilä T, Greco D, Hovatta I. Strong conservation of inbred mouse strain microRNA loci but broad variation in brain microRNAs due to RNA editing and isomiR expression. RNA. 2018;24: 643–655.

33. Engkvist ME, Stratford EW, Lorenz S, Meza-Zepeda LA, Myklebost O, Munthe E. Analysis of the miR-34 family functions in breast cancer reveals annotation error of miR-34b. Sci Rep. 2017;7: 9655.

34. Tan GC, Chan E, Molnar A, Sarkar R, Alexieva D, Isa IM, et al. 5′ isomiR variation is of functional and evolutionary importance. Nucleic Acids Res. 2014;42: 9424–9435.

35. Kume H, Hino K, Galipon J, Ui-Tei K. A-to-I editing in the miRNA seed region regulates target mRNA selection and silencing efficiency. Nucleic Acids Res. 2014;42: 10050–10060.

36. Lukasik A, Wójcikowski M, Zielenkiewicz P. Tools4miRs – one place to gather all the tools for miRNA analysis. Bioinformatics. 2016;32: 2722–2724.

37. Urgese G, Paciello G, Acquaviva A, Ficarra E. isomiR-SEA: an RNA-Seq analysis tool for miRNAs/isomiRs expression level profiling and miRNA-mRNA interaction sites evaluation. BMC Bioinformatics. 2016;17: 148.

38. Knut R, Hailemariam DT, Marcel E, Hannes H, Svenja M, René R, et al. The SeqAn C++ template library for efficient sequence analysis: A resource for programmers. J Biotechnol. 2017;261: 157–168.

39. Barturen G, Rueda A, Hamberg M, Alganza A, Lebron R, Kotsyfakis M, et al. sRNAbench: profiling of small RNAs and its sequence variants in single or multi-species high-throughput experiments. Methods in Next Generation Sequencing. 2014;1. doi:10.2478/mngs-2014-0001

40. Rueda A, Barturen G, Lebrón R, Gómez-Martín C, Alganza Á, Oliver JL, et al. sRNAtoolbox: an integrated collection of small RNA research tools. Nucleic Acids Res. 2015;43: W467–73.

41. Lu Y, Baras AS, Halushka MK. miRge 2.0 for comprehensive analysis of microRNA sequencing data. BMC Bioinformatics. 2018;19: 275.

42. Desvignes T, Batzel P, Sydes J, Frank Eames B, Postlethwait JH. miRNA analysis with Prost! reveals evolutionary conservation of organ-enriched expression and posttranscriptional modifications in three-spined stickleback and zebrafish [Internet]. 2018. doi:10.1101/423533

43. GFF3. In: Sequence Ontology [Internet]. [cited 24 Dec 2018]. Available: https://github.com/The-Sequence-Ontology/Specifications/blob/master/gff3.md

44. miRTop incubator. In: miRTop [Internet]. [cited 24 Dec 2018]. Available: https://github.com/miRTop/incubator/issues

45. Lesurf R, Cotto KC, Wang G, Griffith M, Kasaian K, Jones SJM, et al. ORegAnno 3.0: a community-driven resource for curated regulatory annotation. Nucleic Acids Res. 2015;44: D126–D132.

46. O’Connor BD, Day A, Cain S, Arnaiz O, Sperling L, Stein LD. GMODWeb: a web framework for the generic model organism database. Genome Biol. BioMed Central; 2008;9: R102.

47. Sansone S-A, McQuilton P, Rocca-Serra P, Gonzalez-Beltran A, Izzo M, Lister A, et al. FAIRsharing: working with and for the community to describe and link data standards, repositories and policies [Internet]. 2018. doi:10.1101/245183

48. Ison J, Kalas M, Jonassen I, Bolser D, Uludag M, McWilliam H, et al. EDAM: an ontology of bioinformatics operations, types of data and identifiers, topics and formats. Bioinformatics. 2013;29: 1325–1332.

49. Kozomara A, Griffiths-Jones S. miRBase: annotating high confidence microRNAs using deep sequencing data. Nucleic Acids Res. 2014;42: D68–73.

50. Backes C, Fehlmann T, Kern F, Kehl T, Lenhof H-P, Meese E, et al. miRCarta: a central repository for collecting miRNA candidates. Nucleic Acids Res. 2018;46: D160–D167.

51. Fromm B, Domanska D, Hackenberg M, Mathelier A, Hoye E, Johansen M, et al. MirGeneDB2.0: the curated microRNA Gene Database [Internet]. 2018. doi:10.1101/258749

52. Loher P, Telonis AG, Rigoutsos I. MINTmap: fast and exhaustive profiling of nuclear and mitochondrial tRNA fragments from short RNA-seq data. Sci Rep. 2017;7: 41184.

53. Pliatsika V, Loher P, Telonis AG, Rigoutsos I. MINTbase: a framework for the interactive exploration of mitochondrial and nuclear tRNA fragments. Bioinformatics. 2016;32: 2481–2489.

54. International Human Genome Sequencing Consortium. Finishing the euchromatic sequence of the human genome. - PubMed - NCBI [Internet]. [cited 9 Nov 2018]. Available: https://www.ncbi.nlm.nih.gov/pubmed/15496913

55. Li H, Handsaker B, Wysoker A, Fennell T, Ruan J, Homer N, et al. The Sequence Alignment/Map format and SAMtools. Bioinformatics. 2009;25: 2078–2079.

56. Cock PJA, Antao T, Chang JT, Chapman BA, Cox CJ, Dalke A, et al. Biopython: freely available Python tools for computational molecular biology and bioinformatics. Bioinformatics. 2009;25: 1422–1423.

57. Bofill-De Ros X, Chen K, Chen S, Tesic N, Randjelovic D, Skundric N, et al. QuagmiR: A Cloud-based Application for IsomiR Big Data Analytics. Bioinformatics. 2018; doi:10.1093/bioinformatics/bty843

58. Giraldez MD, Spengler RM, Etheridge A, Godoy PM, Barczak AJ, Srinivasan S, et al. Comprehensive multi-center assessment of small RNA-seq methods for quantitative miRNA profiling. Nat Biotechnol. 2018;36: 746–757.

